# Sustainable production of 2,3,5,6-Tetramethylpyrazine at high titer in engineered *Corynebacterium glutamicum* using statistical design of experiments

**DOI:** 10.1101/2023.10.20.563186

**Authors:** Aparajitha Srinivasan, Kevin Chen-Xiao, Deepanwita Banerjee, Asun Oka, Venkataramana R Pidatala, Aymerick Eudes, Blake A. Simmons, Thomas Eng, Aindrila Mukhopadhyay

## Abstract

**Background:** The industrial amino acid production workhorse, *Corynebacterium glutamicum* naturally produces low levels of 2,3,5,6-tetramethylpyrazine (TMP), a valuable flavor, fragrance and commodity chemical. We have previously reported serendipitous production of TMP in *C. glutamicum* strains. In the present study, we demonstrate TMP production in *C. glutamicum* type strain ATCC13032 via the expression of a heterologous TMP pathway in a defined medium followed by statistical design of experiments to understand the effect of the media composition on TMP production.

**Results:** The *C. glutamicum* strain engineered to overexpress acetolactate synthase and alpha-acetolactate decarboxylase from *Lactococcus lactis* produced ∼0.8 g/L TMP in CGXII minimal medium supplemented with 40 g/L glucose in 24-deep well plates. This engineered strain also demonstrated growth and TMP production when the minimal medium was supplemented with up to 40% (v/v) hydrolysates derived from ionic liquid pretreated sorghum biomass. A screen for improvements in media composition on TMP titer was conducted using fractional factorial design that identified glucose and urea as significant components affecting TMP production. These two components were further optimized using response surface methodology. In the optimized CGXII medium, the engineered strain could produce up to 3.56 g/L TMP (4-fold enhancement in titers and 2-fold enhancement in yield, mol/mol) from 80 g/L glucose and 11.9 g/L urea in shake flask batch cultivation.

**Conclusions:** We engineered the industrially relevant host, *C. glutamicum* for targeted production of TMP by heterologous expression of pathway proteins. We demonstrated the capability of the engineered strain for growth and TMP production utilizing real world carbon streams such as hydrolysates. We further identified glucose and urea as the key minimal media components significantly affecting TMP production using statistical media optimization.

## 1. Introduction

2,3,5,6-Tetramethylpyrazine (TMP), a nitrogen containing heterocyclic aromatic compound, is used widely as a flavor additive in the food industry [1, 2]. Chemically, it can be synthesized using the Maillard reaction [3]. Biologically, TMP production has been reported in bacteria such as *Bacillus sp* [4–12], *Lactococcus lactis* [13, 14]*, Corynebacterium glutamicum* [15–17], *Escherichia coli* [18–20], fungi such as *Saccharomyces cerevisiae* and *Tolypocladium inflatum* [21, 22]. Production of TMP via microbial conversion routes in a more cost-effective and environmentally friendly manner could help to meet its ever-increasing global demand.

The proposed pathway for TMP biosynthesis is depicted in **Figure 1**. TMP is formed from condensation of two molecules of acetoin with ammonia. Acetoin in turn is synthesized from acetolactate via pyruvate decarboxylation [23]. Acetolactate can either undergo spontaneous decarboxylation to form diacetyl which is either reduced to acetoin as reported in *C. glutamicum* [15, 16] or can be decarboxylated enzymatically via alpha-acetolactate decarboxylase (*budA*) to acetoin as in *B. subtilis, B. licheniformis* and *L. lactis* [1]. Acetoin can also be converted to 2,3-butanediol (2,3-BDO) in these microbes. Previous reports have shown that overexpression of the pathway enzymes and/or gene deletions blocking the competing pathways can enhance the carbon flux to TMP [9, 15, 18, 21].

**Figure 1.**
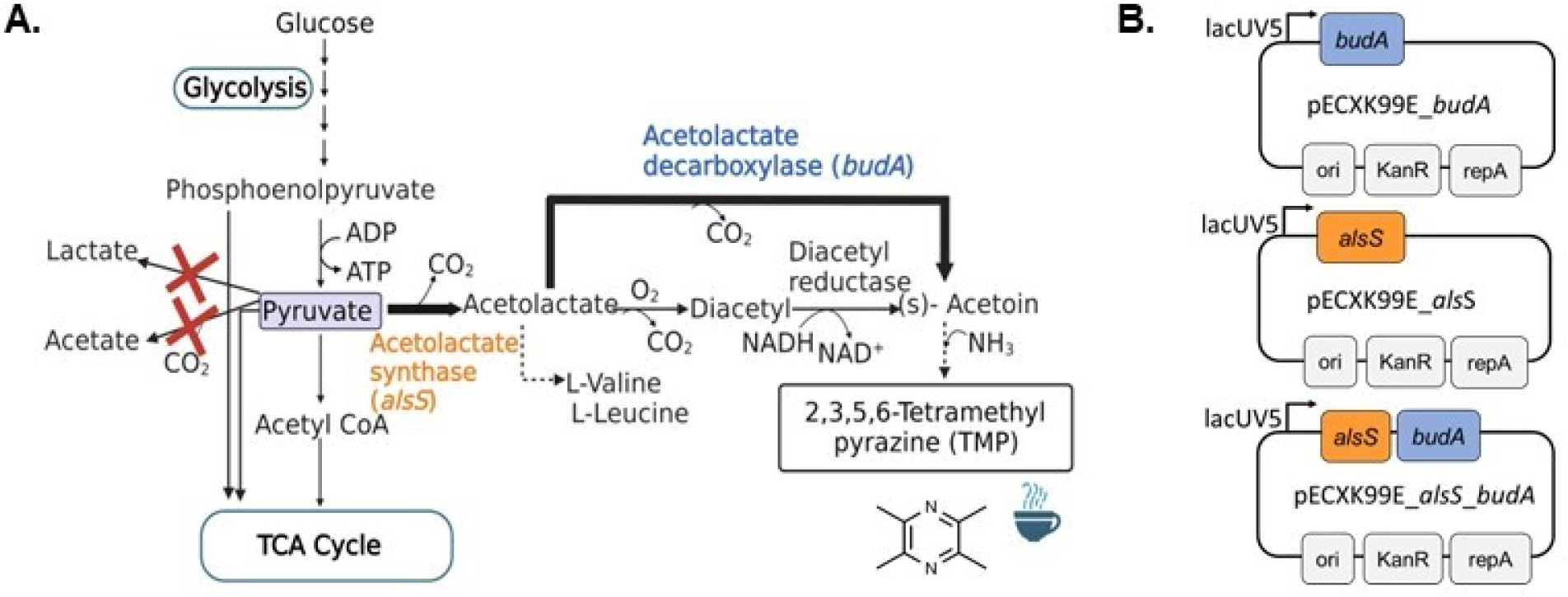
The proposed TMP biosynthesis pathway in *C. glutamicum* and the overexpression constructs employed in this study. A. The overexpressed genes are denoted by bold arrows and the deleted reactions in red. B. The genes *alsS* and *budA* were amplified from either *Bacillus subtilis* (*Bs*) or *Lactococcus lactis* (*Ll*) genome.

*C. glutamicum* is a gram-positive facultative anaerobe that is generally recognized as safe (GRAS). It is an industrially relevant host used widely for bioproduction of amino acids, organic acids, aromatics etc [24, 25]. We previously observed serendipitous production of TMP in *C. glutamicum* engineered for the biofuel precursor, isoprenol [15]. Functional genomics of the engineered strain revealed overexpression of TMP pathway genes amongst others leading to enhanced production of TMP compared to the negligible levels in the wild-type strain [26].

A chemically defined minimal medium which ensures reproducibility and less interference with downstream purifications is often used for bio-based production using microbial hosts. Such a synthetic minimal medium used for routine cultivation of *C. glutamicum* is CGXII [27]. Many studies have explored the effect of change in this medium composition on growth and/or production [18, 24, 25, 28–31]. Despite advances in high throughput-omics and machine learning guided optimization strategies, conventional media optimization strategies remain prominent due to their ease of implementation.

In this study, we hypothesized that engineering *C. glutamicum* to overexpress acetolactate synthase and/or alpha-acetolactate decarboxylase (*als*S*, budA*) would lead to higher TMP titers. To this end, we generated and screened a high TMP producing engineered strain. We then assessed the significance of different components in CGXII minimal medium on growth and TMP production in this engineered *C. glutamicum* strain via statistical design of experiments to obtain a more systematic understanding of these parameters. The engineered *C. glutamicum* strain stably produced ∼0.8 g/L of TMP in CGXII minimal medium supplemented with 40 g/L glucose in 24-deep well plate format. The engineered strain was also able to produce TMP when supplemented with sorghum hydrolysates) for co-utilization of plant biomass precursors. The production was demonstrably optimizable leading to up to a 4-fold increase in TMP titers (up to 3.5 g/L) using statistical tools compared to the control (0.8 g/L) in a microbioreactor.

## 2. Materials and Methods

### 2.1. Chemicals and Reagents

Chemicals and reagents (molecular biology grade or higher) were purchased from Sigma-Aldrich (St. Louis, MO) or as indicated.

### 2.2 Strains and Plasmids

**Table 1** lists the strains and plasmids used in this study and their sequences are available at http://public-registry.jbei.org [32]. Isothermal DNA assembly [33] using the HiFi DNA Assembly Master Mix was used according to manufacturer’s protocols (NEBuilder, New England Biolabs, Ipswich, MA) to assemble pathway gene overexpression plasmid constructs. Oligonucleotides for plasmid assembly and PCR were synthesized by Integrated DNA Technologies, Inc. (San Diego, CA) and listed in **Table S1**.

**Table 1.**
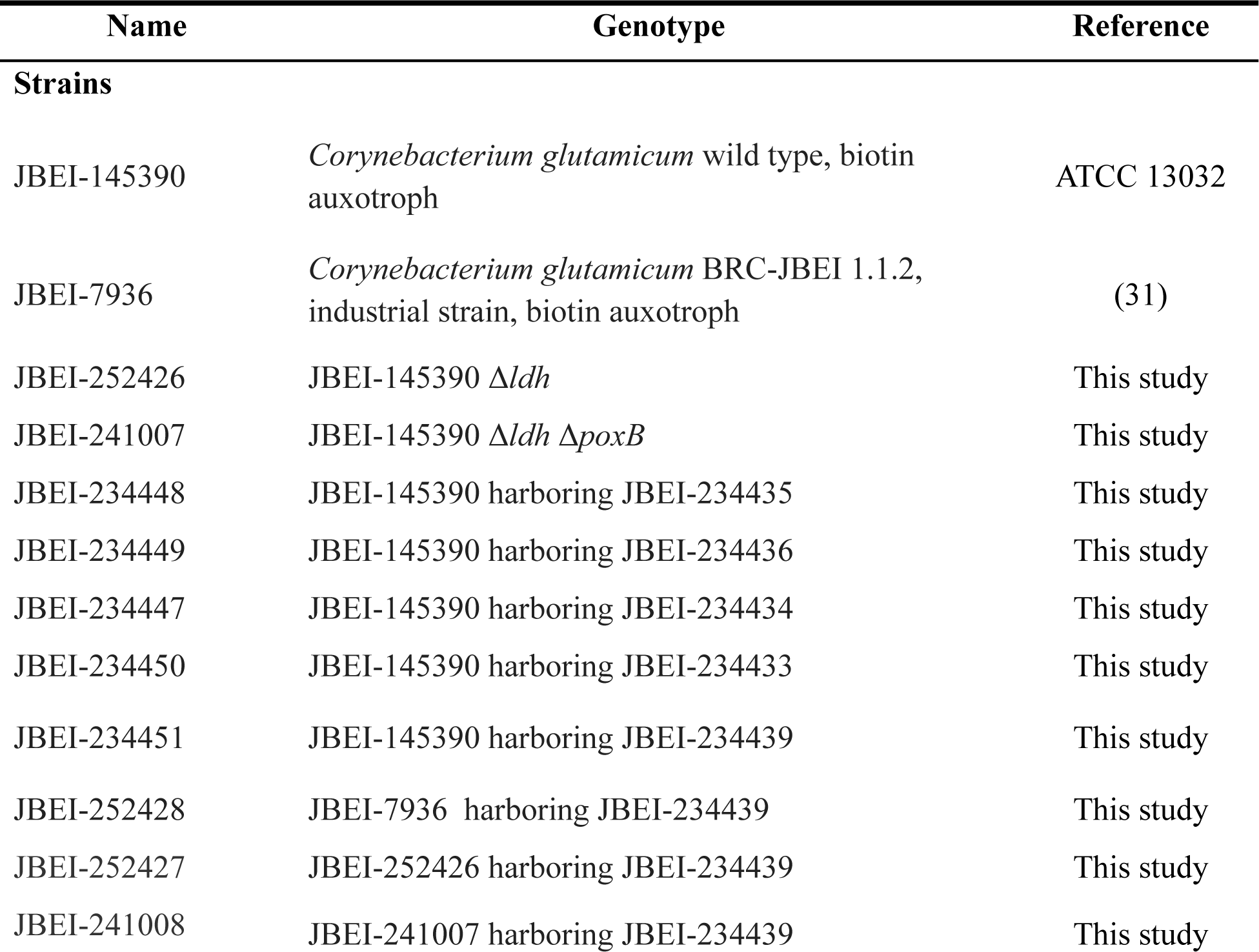
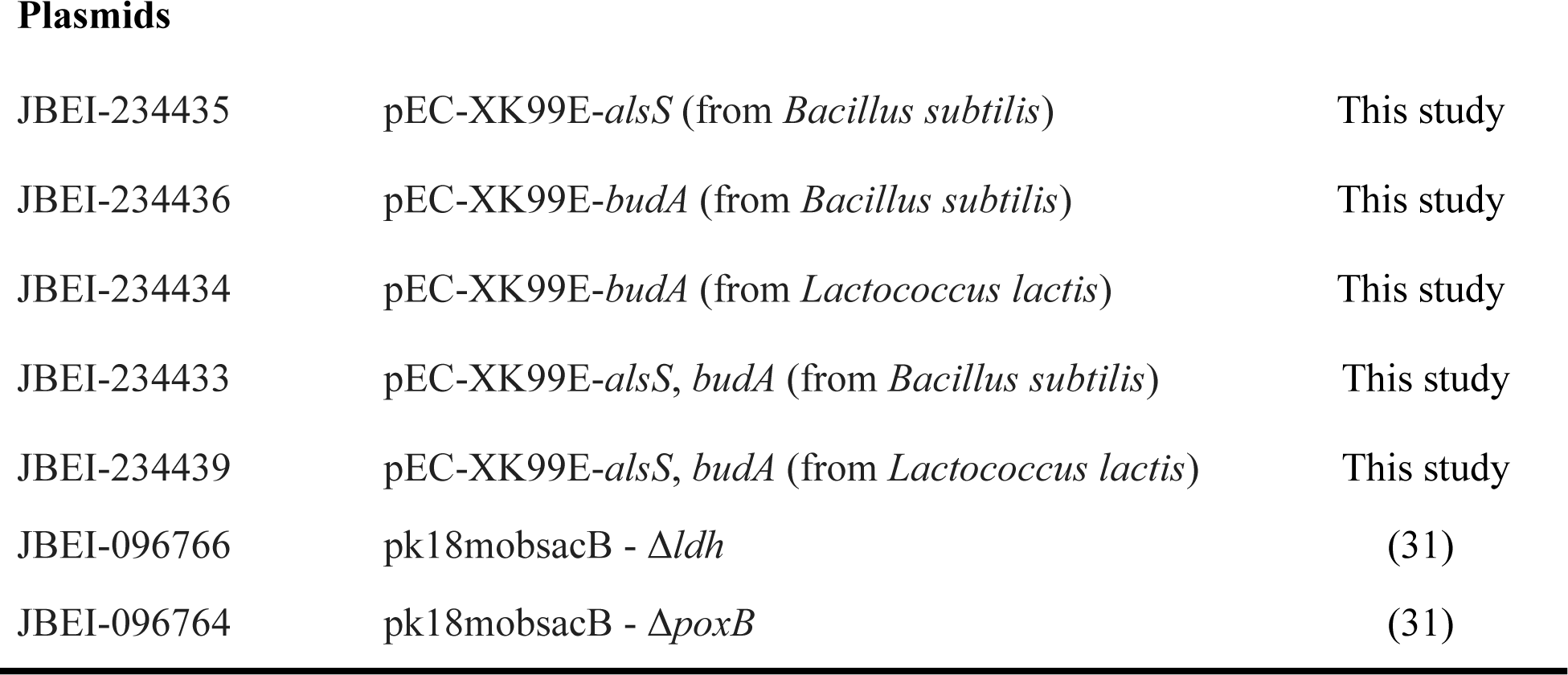
Strains and Plasmids Used in This Study.

| Name | Genotype | Reference |
| --- | --- | --- |
| **Strains** |  |  |
| JBEI-145390 | *Corynebacterium glutamicum* wild type, biotin auxotroph | ATCC 13032 |
| JBEI-7936 | *Corynebacterium glutamicum* BRC-JBEI 1.1.2, industrial strain, biotin auxotroph | (31) |
| JBEI-252426 | JBEI-145390 $\Delta ldh$ | This study |
| JBEI-241007 | JBEI-145390 $\Delta ldh \Delta poxB$ | This study |
| JBEI-234448 | JBEI-145390 harboring JBEI-234435 | This study |
| JBEI-234449 | JBEI-145390 harboring JBEI-234436 | This study |
| JBEI-234447 | JBEI-145390 harboring JBEI-234434 | This study |
| JBEI-234450 | JBEI-145390 harboring JBEI-234433 | This study |
| JBEI-234451 | JBEI-145390 harboring JBEI-234439 | This study |
| JBEI-252428 | JBEI-7936 harboring JBEI-234439 | This study |
| JBEI-252427 | JBEI-252426 harboring JBEI-234439 | This study |
| JBEI-241008 | JBEI-241007 harboring JBEI-234439 | This study |
| Plasmids |  |  |
| JBEI-234435 | pEC-XK99E-*alsS* (from *Bacillus subtilis*) | This study |
| JBEI-234436 | pEC-XK99E-*budA* (from *Bacillus subtilis*) | This study |
| JBEI-234434 | pEC-XK99E-*budA* (from *Lactococcus lactis*) | This study |
| JBEI-234433 | pEC-XK99E-*alsS*, *budA* (from *Bacillus subtilis*) | This study |
| JBEI-234439 | pEC-XK99E-*alsS*, *budA* (from *Lactococcus lactis*) | This study |
| JBEI-096766 | pk18mobsacB - $\Delta ldh$ | (31) |
| JBEI-096764 | pk18mobsacB - $\Delta poxB$ | (31) |

For error-free PCR, Q5 High-Fidelity DNA Polymerase (New England Biolabs, Ipswich, MA) was used. Plasmids were initially transformed into chemically competent *E. coli* XL-1 Blue (New England Biolabs) and all sequences were confirmed by colony PCR and whole plasmid sequencing (Primordium Labs, Arcadia, CA) followed by electroporation into *C. glutamicum* ATCC13032 as described previously [31]. For the TMP pathway engineering, alpha-acetolactate decarboxylase (*budA*) and acetolactate synthase (*alsS*) genes from *L. lactis* (ATCC 19257) and *B. subtilis* (ATCC 6051) were overexpressed either individually or in combination using the *C. glutamicum* shuttle expression vector pEC-XK99E.

### 2.3 Production of TMP from Engineered *C. glutamicum*

A single colony was picked from LB agar plates containing kanamycin (50 mg/L) following standard laboratory procedures. Colonies were inoculated and grown in 5 mL LB medium (with kanamycin) at 30 °C on a rotary shaker at 200 rpm overnight. Cultures were then adapted by back diluting saturated cultures in 5 mL CGXII minimal medium (28) for 24 h twice prior to starting production runs as described previously (15).

For production runs in 24-deep well plates, 2 mL of culture medium (CGXII with 40 g/L glucose) was used per well with a starting OD_600_ (optical density at 600 nm) of 0.1. Deep well plates were incubated in a Multitron incubator (Infors HT, Switzerland) with a 3 mm orbital shaking platform shaken at 999 rpm (Bottmingen, Switzerland). The production pathway was induced with 0.5 mM IPTG after inoculation. The runs were monitored for periodic OD_600_ measurements, residual sugars and TMP accumulation at 24 h, 48 h and 72 h.

### 2.4 Production of TMP from Plant Biomass Hydrolysates

CGXII minimal medium was supplemented with hydrolysates obtained from sorghum biomass to test the ability of the engineered high TMP producing strain to utilize real world carbon streams treated with ionic liquids. Sorghum biomass (stems and leaves without panicles) from field-grown wild-type and genetically engineered plants that accumulate high amounts of 4-hydroxybenzoic acid (4-HBA) was obtained as previously described (34). Moreover, ensiled sorghum biomass was obtained from a commercial silage pit on a dairy farm in the southern part of the San Joaquin Valley, California. The hydrolysates were generated as described previously (35), briefly, biomass pretreatment was carried out at a solid loading of 15 wt.% in a one-pot configuration using a 1 L Parr reactor (Parr Instrument Company, model: 4555-58, Moline, IL, USA), 10 wt.% cholinium lysinate [Ch][Lys] and 75 wt.% water. Typically, 60 g of biomass was mixed thoroughly with 40 g of [Ch][Lys] and 300 g of water followed by heating at 140 °C for 3h. Post pretreatment, 10 M HCl was added to adjust the pH of the biomass slurry to 5. Subsequently, a commercial enzyme mixture, Cellic CTec3 and HTec3 (9:1 v/v) (Novozymes North America, Franklinton, NC, USA), was added to the biomass slurry at a concentration of 10 mg enzyme/g biomass to carry out saccharification at 50 °C for 72 h at 48 rpm in the same Parr vessel. Resulting hydrolysate was centrifuged at 8000 rpm for 20 min and the supernatant was filtered using 0.45 µM and 0.22 µM rapid flow filters (Nalgene, USA). The hydrolysate pH was adjusted to 7.4 with 1 N NaOH. Hydrolysates were then added making up to 20% v/v or 40% v/v of the total sterile 1X CGXII media (to a total of 40 g/L glucose by supplementation of pure glucose). The production runs were carried out on 24-deep well plates as described in section 2.3 with the supplemented hydrolysates. OD_600_, TMP (g/L) and residual sugars were measured at 24 h, 48 h and 72 h.

### 2.5 Calculation of Maximum Theoretical Yield of TMP

The maximum theoretical yield of TMP was calculated by using a genome-scale metabolic model (GSMM) of *C*. *glutamicum*, iCGB21FR (36). Reactions (1 - 3) for diacetyl, acetoin, TMP formation and TMP exchange were added to the GSMM for facilitating *in silico* TMP production.

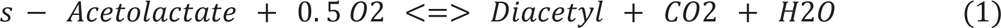

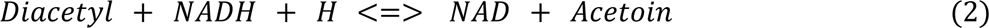

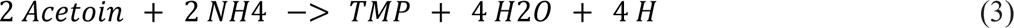

Using flux balance analysis (FBA) (37) and a fixed constraint of 10 mmol/g dry cell weight/h for the glucose uptake, the TMP formation was set as the objective to be maximized in order to calculate the maximum theoretical yield (MTY) of TMP for *C*. *glutamicum*. FBA analysis was conducted using the COBRA toolbox (38) for MATLAB (version R2020a, http://www.mathworks.com) and the solver Gurobi (version 6.0, http://www.gurobi.com).

### 2.6 Selection of Significant Media Components Using Statistical Design of Experiments

Design of experiments was used for testing the effect of CGXII minimal medium components on growth and TMP production. The statistical design and analysis was carried out using the Design-Expert software (Stat-Ease, USA) For the initial fractional factorial design (FFD) to screen significant components in the CGXII minimal media, ten of the media components were evaluated. Each component was examined at two levels, a high (+) and a low (−) level of concentration (**Table 2**) in a resolution IV screening design leading to 32 runs (**Table S2**). It is to be noted that in the resolution IV designs, typically the main effects are not confounded with other main effects or two-factor interactions but aliased with three-factor interactions. Also, certain two-factor interactions could be confounded with one other two-factor or more three-factor interactions. The experimental responses measured as OD_600_ and TMP titers (g/L) in each of these 32 experiments were subjected to statistical analysis. The Design Expert software was used to assess the significance of each component based on the first-order model assumption using equation (4),

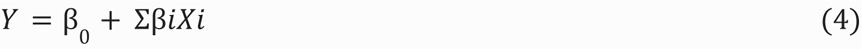

**Table 2.** Concentrations of the CGXII Minimal Media Components Used for Fractional Factorial Design.

| Factor | Component | Levels |  |
| --- | --- | --- | --- |
|  |  | -1 | +1 |
| A | Glucose | 20 | 60 |
| B | Ammonium sulfate (NH<sub>4</sub>)<sub>2</sub>SO<sub>4</sub> | 10 | 40 |
| C | Urea | 2.5 | 10 |
| D | Potassium phosphate monobasic (KH<sub>2</sub>PO<sub>4</sub>) | 0.5 | 2.0 |
| E | Potassium phosphate dibasic (K<sub>2</sub>HPO<sub>4</sub>) | 0.5 | 2.0 |
| F | Magnesium sulfate (MgSO<sub>4</sub>·7H<sub>2</sub>O) | 0.125 | 0.5 |
| G | Trace Elements | 0.5X | 2X |
| H | Biotin | 0.0001 | 0.0004 |
| I | Protocatechuate (PCA) | 0.015 | 0.06 |
| J | Calcium chloride (CaCl<sub>2</sub>) | 0.005 | 0.02 |

All concentrations in g/L except for Trace elements which is depicted in fold (X)

where *Y* is the response predicted, β_0_ is the intercept, *βi* is the linear coefficient or the slope indicating the magnitude of change expected in *Y* when there is one unit change in the medium component *Xi*. Media components with a low *p* value (*p* < 0.05) indicating a significant effect on the responses were selected for subsequent optimization.

Response Surface Methodology (RSM) was then used to determine the optimum concentrations of the significant components, while the rest of the medium components were kept constant. The Design-Expert software was used to generate a 13-run central composite design (CCD) for this purpose from the two significant factors obtained from FFD screening with 2k i.e., 4 full-factorial points (where *k* is the number of components), 5 center points (replicates), and 2 axial points for each component (**Table 3**). Replication of design points provides an estimation of pure error in the design.

**Table 3.** Concentrations of each selected media component at different levels of the Central Composite Design.

| Factor | Component | Unit | Levels |  |  |  |  |
| --- | --- | --- | --- | --- | --- | --- | --- |
| | | | - 1 | 0 | + 1 | - $\alpha$ | + $\alpha$ |
| A | Glucose | (g/L) | 60 | 70 | 80 | 55.86 | 84.14 |
| B | Urea | (g/L) | 10 | 15 | 20 | 7.93 | 22.07 |

Experiments were carried out on 48-well flower plates in a microbioreactor (BioLector Pro, M2P labs, Germany). For runs in the microbioreactor format, cells (inoculated to OD_600_ = 0.1 and induced with 0.5 mM IPTG) were grown in 48-well flower plate (M2P labs, Germany) containing 1 mL medium in each well with antibiotic, sealed with gas permeable membrane (M2P labs, Germany) and shaken at 1200 rpm at 30 °C. The runs were then analyzed for cell density, final residual sugar and TMP at 48 h.

### 2.7 Analytical Methods for Metabolite Quantitation

For metabolite quantification, 100 μL of cell culture medium was combined with 100 μL of ethyl acetate containing n-butanol (30 mg/L) as an internal standard and analyzed as described previously (15). Briefly, post extraction with ethyl acetate, 100 μL of the ethyl acetate phase was transferred into a GC vial with insert and 1 μL was analyzed using GC 8890 (Agilent Technologies, USA) equipped with a flame ionization detector (FID) and a DB-WAX capillary column (Agilent Technologies, USA) for quantification. The temperatures of the injector and detector were 250 and 300 °C, respectively. Helium was used as the carrier gas (2.2 mL/min) and the injection volume was 1 μL in splitless mode. The data was collected and analyzed using OpenLab software (Agilent Technologies, USA). Analytical grade standards were purchased from Sigma-Aldrich (St. Louis, MO) and used to calculate analyte concentrations and confirm peaks of TMP and acetoin. Residual sugars and organic acids were measured using a HPLC 1100 (Agilent Technologies, USA) equipped with an Aminex 75H column with 4 mM sulfuric acid as the mobile phase at 0.6 mL/min and analyzed using a refractive index detector (RID).

## 3. Results and Discussion

### 3.1 Over-expression of *alsS* and *budA* from *L. lactis* Enhances TMP Production in Engineered *C. glutamicum*

We overexpressed TMP pathway enzymes (encoded by *alsS* and *budA*) from *B. subtilis* and *L. lactis* under the control of IPTG inducible promoter using the shuttle expression vector pEC-XK99E in *C. glutamicum* ATCC13032 and tested their effect (individually and in combination) on growth and TMP production. The engineered strain overexpressing both the enzymes from *L. lactis* resulted in a high TMP titer of 898 ± 22 mg/L at 48 h (**Figure 2a**). GC-FID analysis indicated that in the highest TMP producer, acetoin accumulates minimally (<100 mg/L at 24 h and 48 h) whilst TMP titers increase concomitantly (**Figure 2b**). On the other hand, > 0.7 g/L acetoin accumulation was observed at 48 h in the strain(s) overexpressing *budA* from either *B. subtilis* or *L. lactis -* the highest titer (3.81 g/L) observed in the strain overexpressing *budA_Bs_* (**Figure 2b**). The rate of formation of acetoin in these strains could be greater than the rate of spontaneous conversion (dependent on the availability of ammonia) to TMP in the sampled time-points (13). We did not test the enantiomeric form of acetoin produced in these reactions. The TMP formation could also be hindered if the acetoin formed is not the appropriate enantiomeric substrate needed for condensation to TMP. Glucose was completely consumed by 48 h across all the different engineered strains (data not shown) and considering the accumulation of TMP and its precursors in the late exponential/stationary phase, the production does not seem to impact growth (**Figure 2c**) and glucose consumption significantly. We also did not observe any significant accumulation of overflow metabolites such as lactate or acetate in these transformants. Since the TMP titer was maximum at 48 h, in the remaining experiments we focused on harvesting the cells and reporting titers at this time point. Preliminary evaluation showed that overexpression of *alsS_Ll_* alone also led to higher levels of TMP (data not shown). Nevertheless, we chose the engineered strain with the *alsS*_Ll_-*budA*_Ll_ overexpression, which we hypothesized to be a more reliable producer of the acetoin precursor.

**Figure 2.**
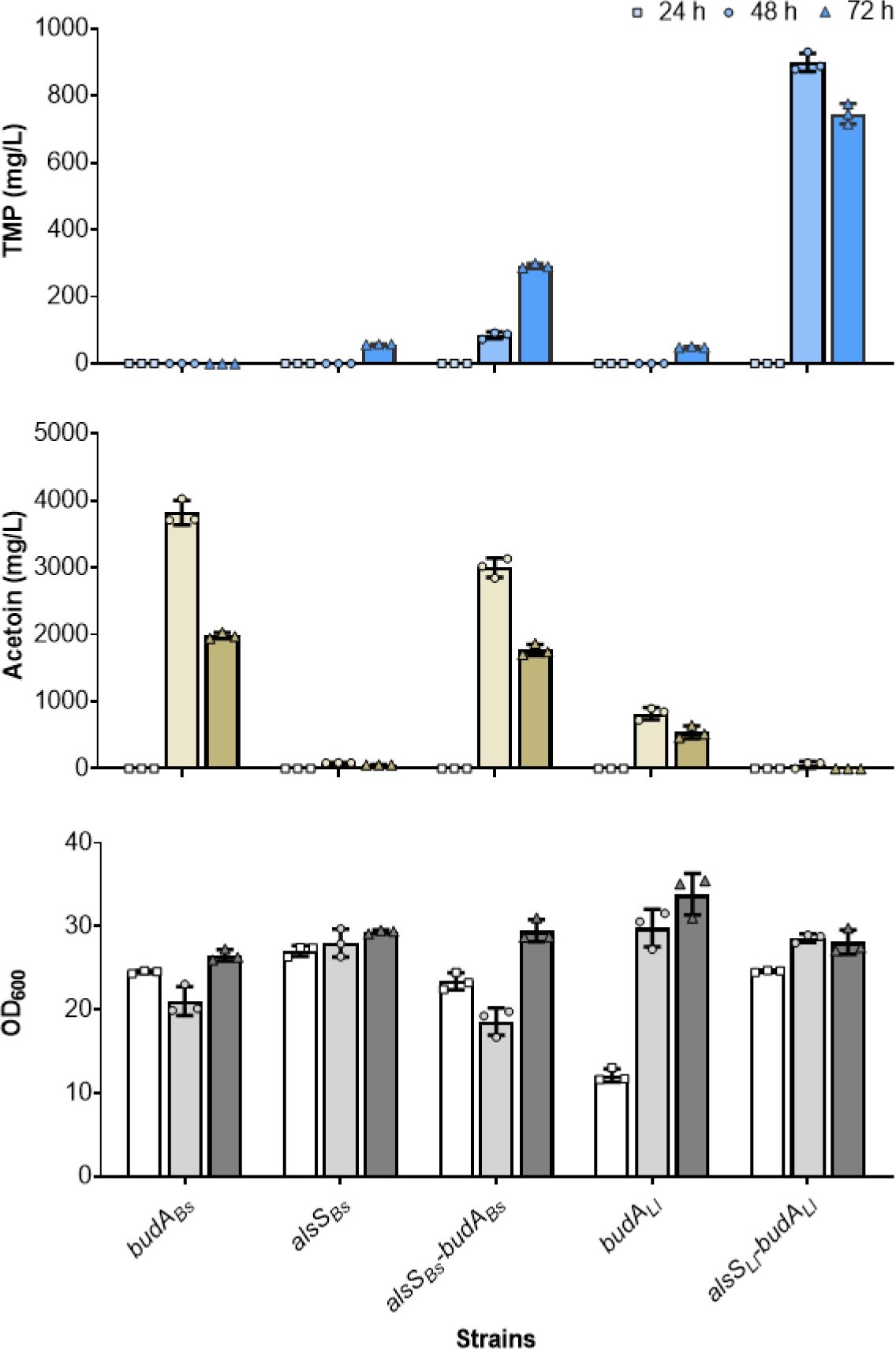
TMP, acetoin titers (mg/L) and cell density (OD_600_) profiles in the various engineered strains. *C. glutamicum* ATCC13032 was transformed with pEC-XK99E overexpressing the respective single (*alsS*_Bs_, *budA*_Bs_, *budA_Ll_*) and double gene (*alsS*_Bs_*-budA*_Bs_, *alsS*_Ll_-*budA*_Ll_) cassettes derived from either *B. subtilis* (Bs) or *L. lactis* (Ll). Strains were grown in 2 mL CGXII minimal medium with 50 mg/L kanamycin and 4% glucose in 24-deep well plates. Samples were harvested at 24 h, 48 h and 72 h. Data represented as average ± SD (n = 3 replicates).

Since pyruvate is the main precursor for TMP, we generated deletions (Δ*ldh* and Δ*ldh* Δ*pox*B) in the parent strain (ATCC13032) to reduce flux diversion to competing metabolites, lactate and acetate as demonstrated in (39, 19), via allelic gene exchange. However, we did not see any significant differences in TMP titers upon deletion of *ldh* or the combined deletion of *ldh* and *poxB* (**Figure 3**). This observation could be specific to the tested media and culture conditions as the flux redirection to overflow (lactate and acetate formation) in the parent strain was much less (11 and 23 mg/L respectively) to start with.

**Figure 3.**
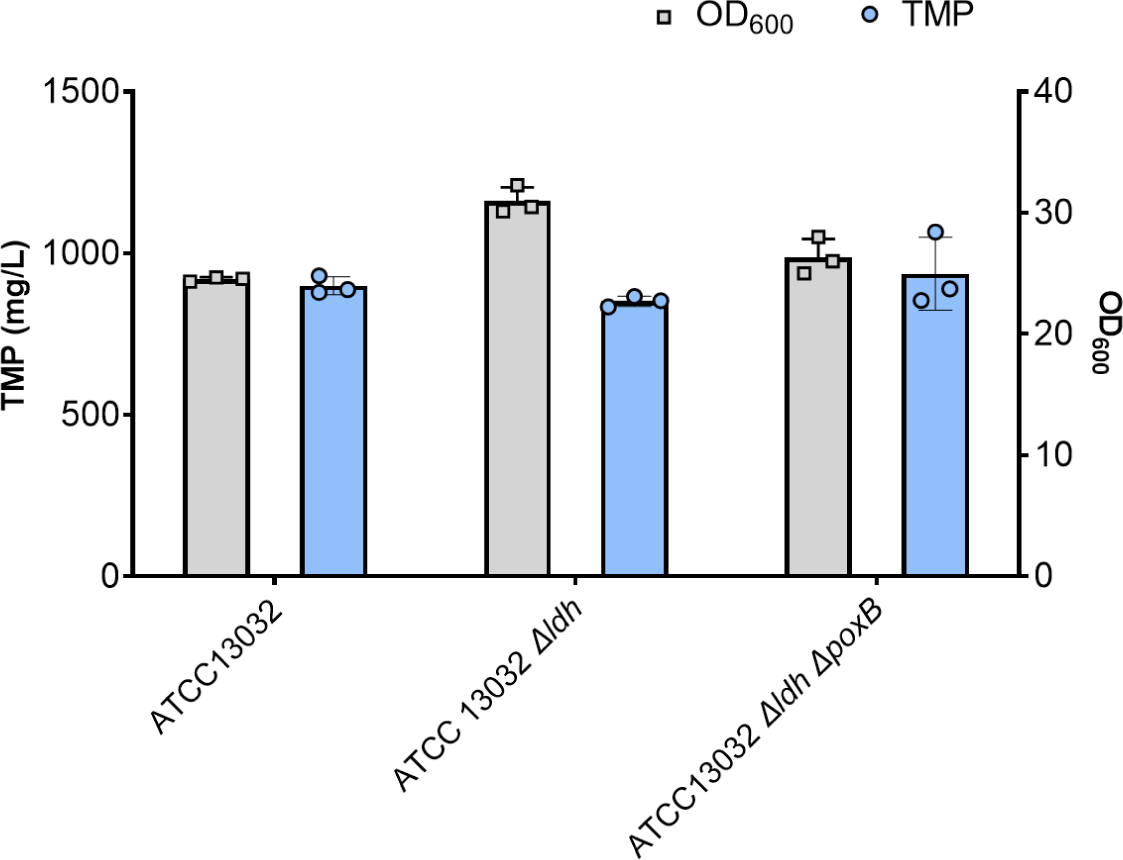
Effect of inhibition of competing pyruvate metabolism on TMP titers (mg/L) in engineered *C. glutamicum*. The strains (ATCC13032 type strain, ATCC13032 Δ*ldh* and ATCC13032 Δ*ldh* Δ*poxB*) overexpressing *alsS_Ll_*-*budA_Ll_* were grown in 2 mL CGXII minimal medium with 4% glucose, 50 mg/L kanamycin in 24-deep well plates and induced with 0.5 mM IPTG. Samples were harvested after 48 h. Data represented as average ± SD (n = 3 replicates).

### 3.2 The Engineered *C. glutamicum* Type Strain Converted Acetoin to TMP More Efficiently than the Strain BRC-JBEI 1.1.2

In a previous study, we observed serendipitous production of TMP in the strain *C. glutamicum* BRC-JBEI 1.1.2 engineered for isoprenol upon treatment with 10% cholinium lysinate ([Ch][Lys]) and under fed-batch bioreactor cultivation (15). The BRC-JBEI 1.1.2 strain is 99.9987% identical to *C. glutamicum* SCgG1 and SCgG2 and only 89% similar to the type strain ATCC13032 with unique genes allowing for a comparative analysis for production. Interestingly, we saw ∼5-fold lower TMP titers when we transformed this strain with the plasmid overexpressing *alsS_Ll_*-*budA_Ll_*(JBEI-234439) and without significant changes in acetoin production compared to the WT strain (**Figure 4a**). We also observed acetate accumulating up to 1 g/L in the same strain at 48 h. Since the last step in the TMP formation is spontaneous and dependent on ammonia, we hypothesize that the residual carbon (including the excessive acetate overflow as seen in **Figure 4b**) and nitrogen may affect the formation differentially in these two different *C. glutamicum* base strains engineered to make TMP. Therefore, we focused on the engineered type strain (ATCC13032 overexpressing *alsS_Ll_*-*budA_Ll_*) for further optimization due to its superior performance.

**Figure 4:**
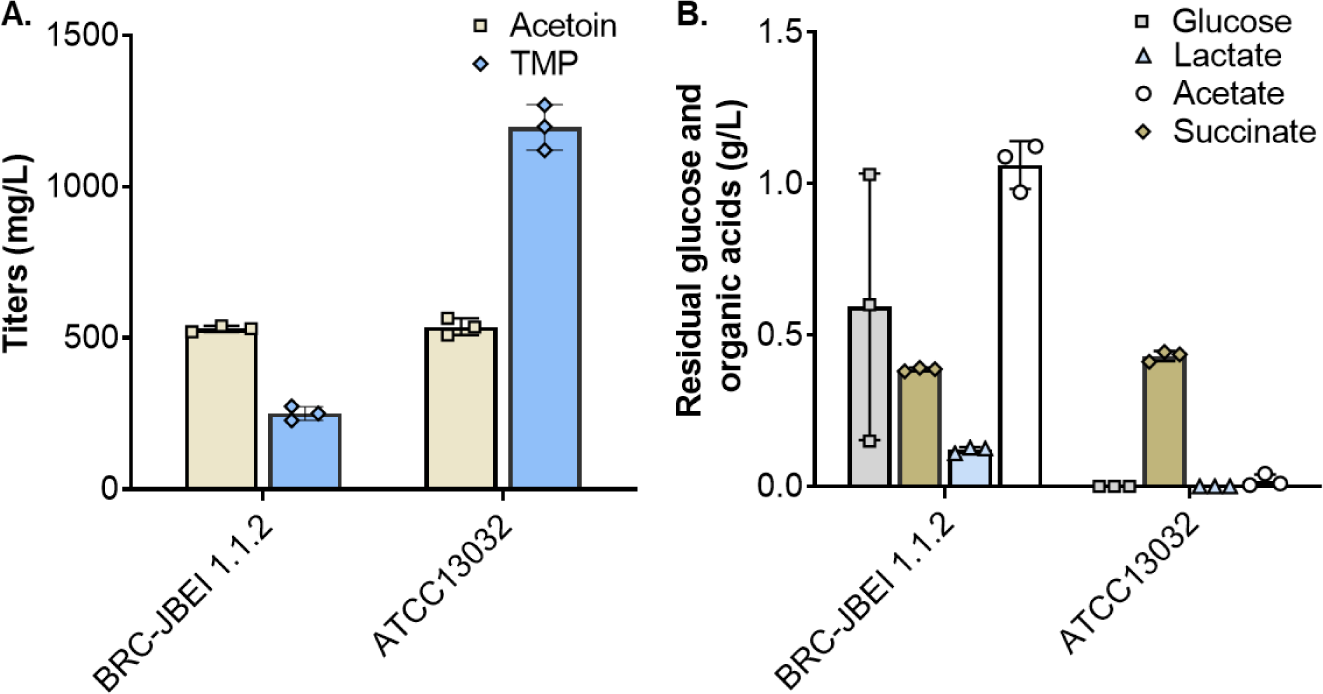
TMP production in the engineered *C. glutamicum* BRC-JBEI 1.1.2 and ATCC13032 strains overexpressing *alsS_Ll_*-*budA_Ll_*. Production of TMP (mg/L) and acetoin (mg/L) (a). Residual glucose and organic acid profiles (g/L) in the spent medium of the two engineered strains analyzed using HPLC after 48 h (b). The strains were grown in 24-deep well plates containing 2 mL CGXII minimal medium with 4% glucose, 50 mg/L kanamycin and induced with 0.5 mM IPTG. Data represented as average ± SD (n = 3 replicates).

### 3.3 Growth and TMP Production of the Engineered C. *glutamicum* on Lignocellulose Derived Carbon Streams

producer from the present study to utilize hydrolysates from ionic liquid ([Ch][Lys]) treated ensiled or 4-HBA enriched sorghum biomass. The engineered strain showed no growth defects when grown with up to 40 % (v/v) hydrolysate supplemented media and produced TMP in the range of 500-660 mg/L at 48 h upon supplementation with 20% (v/v) sorghum biomass hydrolysates which is slightly lower than the pure glucose supplemented minimal medium (**Figure 5**).

**Figure 5.**
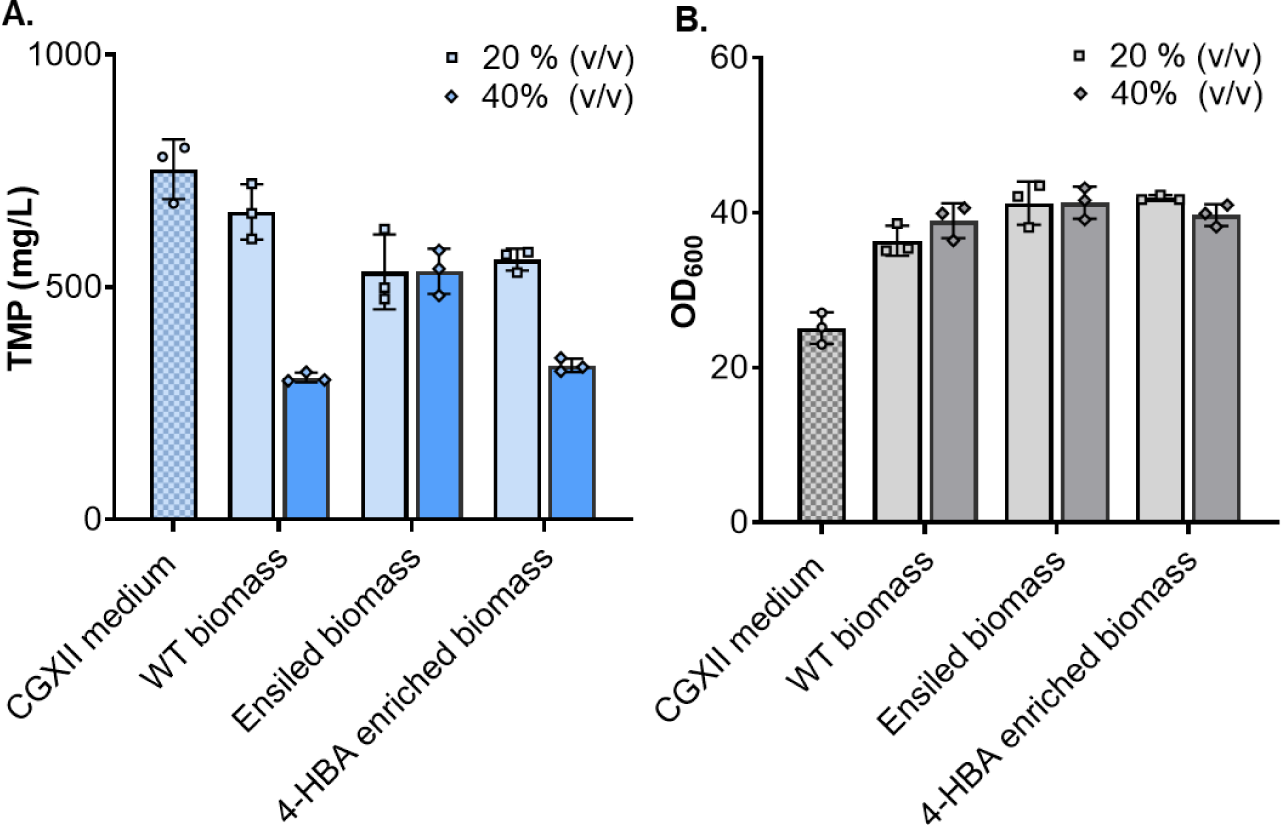
Growth and production of TMP with sorghum hydrolysate supplementation. OD_600_ and TMP titer (mg/L) in the TMP producer strain overexpressing *alsS_Ll_*-*budA_Ll_* on CGXII minimal medium supplemented with 20% (v/v) and 40% (v/v) hydrolysates from wildtype (WT), 4-HBA enriched, or ensiled sorghum biomass. The strain was grown in 24-deep well plates and induced with 0.5 mM IPTG and samples harvested at 48 h. Data represented as average ± SD (n = 3 replicates).

As the hydrolysate volume was increased to 40%, the WT and 4-HBA biomass supplementation exhibited further decrease in TMP titers compared to 20% (v/v) whereas the ensiled biomass could still maintain similar titers as that of 20% (v/v). On the other hand, it was interesting to note the significantly increased accumulation of acetoin (up to 2 g/L) upon supplementation with 20% (v/v) WT and ensiled hydrolysates compared to the minimal medium (**Figure S1**). Despite low titers of TMP in the complex hydrolysate supplemented medium, significant acetoin accumulation and retention of growth (in the presence of possible inhibitory compounds) leaves room for improvement. As the hydrolysates possibly contain various other plant biomass derived components and residues carried over from the pretreatment steps which might affect the metabolic flux to TMP (or) alter the carbon to nitrogen ratio for enhanced production of TMP, we envisage sustainable production with further process optimization.

### 3.4 Glucose and Urea are Significant Minimal Medium Components Affecting TMP Production in Engineered *C. glutamicum*

As medium composition impacts growth and productivity of *C. glutamicum* (29–31), we hypothesized that optimizing medium components could result in improved production. We predicted a maximum theoretical yield (MTY) of 0.58 mol TMP per mol of glucose (i.e. 0.44 g TMP/g glucose or 17.66 g/L TMP from 40 g/L of glucose) using the genome scale metabolic model, iCGB21FR. Experimentally, in CGXII minimal medium, the highest TMP producing strain resulted in 0.03 mol TMP/mol glucose, i.e. only 5.2 % of the MTY, indicating considerable room for improvement. In a previous report we varied the carbon:nitrogen ratio and identified local maxima that improved isoprenol titer (31). However, CGXII medium consists of >12 components, which could generate a larger search space for unanticipated determinants of high TMP titer. To identify significant medium components affecting production systematically, we used a design of experiments approach (29). Given the large number of components in the CGXII medium, we chose to screen the main components affecting growth and/or TMP production using FFD that helps to identify the significant components utilizing limited experiment runs and resources. To prevent media and product evaporation during tests and ensure high mass transfer by effective mixing and scalability, the cells were grown in a microbioreactor in 48-flower well microtiter plates. We retained usage of the standard 0.5 mM IPTG for induction as it did not significantly affect titers in the range of 0.125 - 1.5 mM and a fill volume of 1 mL for the flower plates for further experiments based on preliminary optimizations (**Figure S2 and Figure S3**).

The biomass and TMP titers obtained after 48 h in each of the experiment runs (**Table S2**) were subjected to analysis by the Design-Expert software (Stat-Ease, USA). Our results indicated that glucose (A, the major carbon source) and its possible interaction (AC) with urea (C, one of the nitrogen sources) affected cell density (represented by OD_600_) whereas glucose (A), urea (C) and their possible interaction (AC) significantly affected TMP production (**Table 4**). The other medium components were not significant in their respective tested range. It is to be noted that in this resolution IV screening design used, the interactive effect (AC) is confounded by another two-factor interaction (biotin with calcium chloride as well as other higher order interactions). We assume that the AC interaction effect is likely significant since the main effects (A and C) involved are also significant according to the heredity principle. High TMP titers (2 to 3 g/L) with high cell densities were obtained in runs 3, 11, 15, 17, 24 and 27 with 60 g/L glucose and 10 g/L urea (highest in the range tested). Similarly, cell density and TMP accumulation were relatively low when these medium components were low (where mostly TMP was not detected). Their significance is clear from the *p*-value in the ANOVA for the selected factorial model indicating these components as significant (*p*<0.05) (**Table 4**).

**Table 4.** *p*-values from the ANOVA for Screening of Significant Medium Components Affecting Cell Density and TMP Production in the Fractional Factorial Design (FFD)

| Factor | *p*-value from ANOVA for Selected Factorial Model |  |
| --- | --- | --- |
|  | Cell Density (OD<sub>600</sub>) | TMP (mg/L) |
| A- Glucose | < 0.0001 | < 0.0001 |
| C- Urea | 0.1290 | < 0.0001 |
| AC- Interaction of A and C | <0.0001 | < 0.0001 |

Tolerance to and utilization of high glucose concentrations up to 140 g/L has been demonstrated in batch cultivations of *C. glutamicum*. Given the positive impact on TMP production with increasing glucose concentration, we explored higher glucose ranges for the second level of optimization (up to 80 g/L glucose) in the CGXII medium. Our results also concur with established studies (29) that urea is preferred to ammonium sulfate and affects growth and product formation significantly.

### 3.5 Optimizing the Significant Minimal Medium Components using Response Surface Methodology (RSM)

Optimum concentrations of glucose and urea were determined using RSM, while the rest of the medium components were kept at constant levels (reduced to half the concentrations in CGXII medium except for MOPS). A 13-run central composite design (CCD), developed using the Design-Expert software (Stat-Ease, USA), was used to optimize the concentrations of the two factors. The experimental recipe and response in CCD for individual runs is given in **Table 5**.

**Table 5.** The Central Composite Design (CCD) and Responses Obtained.

| Run | Factor 1:<br>Glucose (g/L) | Factor 2: Urea<br>(g/L) | Cell Density<br>(OD<sub>600</sub>) | TMP (mg/L) |
| --- | --- | --- | --- | --- |
| 1 | 70 | 7.92 | 31.7 | 2402.679 |
| 2 | 70 | 15 | 35.6 | 2519.067 |
| 3 | 60 | 10 | 30.25 | 2161.951 |
| 4 | 80 | 10 | 32.5 | 3220.721 |
| 5 | 70 | 15 | 36.5 | 2326.03 |
| 6 | 55.86 | 15 | 39.8 | 573.992 |
| 7 | 70 | 22.07 | 43.9 | 19.015 |
| 8 | 80 | 20 | 32.2 | 1813.542 |
| 9 | 70 | 15 | 38.5 | 2275.241 |
| 10 | 70 | 15 | 30 | 2624.174 |
| 11 | 60 | 20 | 40.9 | 35.444 |
| 12 | 70 | 15 | 34.5 | 2355.589 |
| 13 | 84.14 | 15 | 34.1 | 2880.99 |

The data on cell density (OD_600_) and TMP production from RSM were subjected to an analysis of glucose and urea tested, indicating the importance of this specific carbon to nitrogen ratio on TMP production and the importance of design of experiments as demonstrated previously (18). We were able to fit the TMP production (mg/L) response at 48 h using a quadratic model (the highest order polynomial where the additional terms are significant, and the model was not aliased) to quantify the individual and interactive effects of glucose (A) and urea (B) as follows

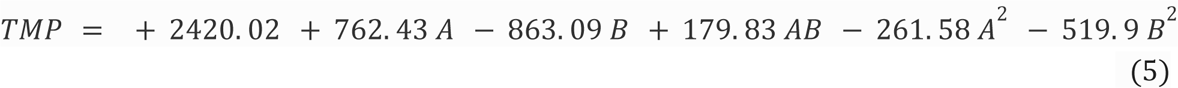

Contour plot (2D) generated by the software was used to visualize the interactive effect of the glucose and urea on TMP production and to find the optimized concentrations of the same (**Figure 6**). The statistical significance of the model equation was confirmed by ANOVA (**Table S2**).

**Figure 6.**
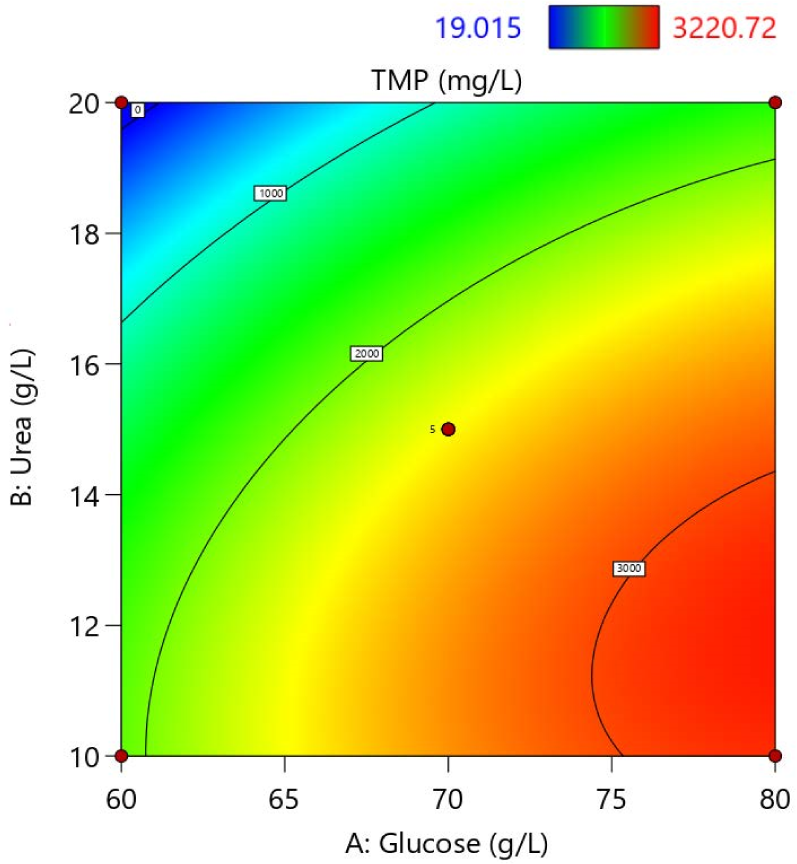
Contour plot (2D) showing the significance and interaction of glucose and urea on TMP production. Glucose (Component A), the main carbon source and the significant nitrogen source, urea (Component B) and their effect on the response, TMP (mg/L). The values within the white squares are experimentally observed values of TMP at these design levels. A total of 4 design points (red circles), 4 corner points and 1 center point (replicated 5 times) was used in the design.

Based on the model, an experiment was run using the optimum values for both glucose and urea as given by the response optimization to confirm that the predicted values of the TMP were obtained. The model predicted optimum (80 g/L glucose and 11.9 g/L urea) resulted in 3.56 g/L TMP (**Figure 7**) reproducibly in two different growth formats - microbioreactor and batch shake flasks.

**Figure 7.**
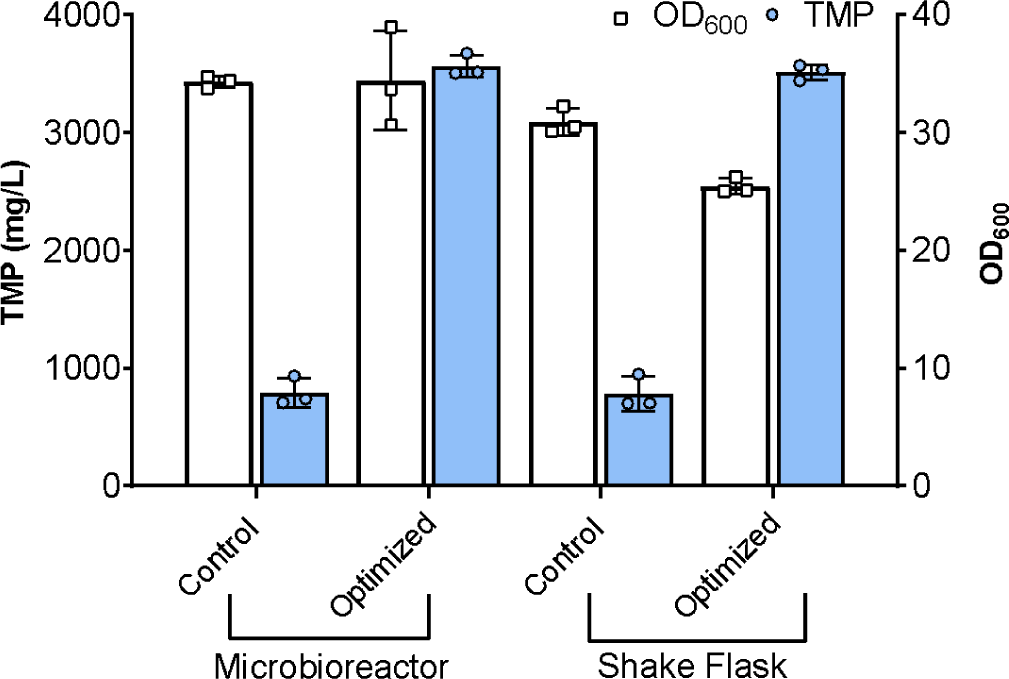
TMP (mg/L) production in the engineered *C. glutamicum* strain grown in the control and the optimized CGXII minimal medium. The strain overexpressing *alsS*_Ll_-*budA*_Ll_ was grown in 48-well flower plates in a microbioreactor as well as in 50 mL of medium in 250 mL baffled shake flasks and induced with 0.5 mM IPTG. Samples were harvested at 48 h. Data represented as average ± SD (n = 3 replicates).

In the optimized medium all the other CGXII media components (except for the MOPS buffering component) were reduced to half their initial value (**Table 6**). Interestingly, there was an increased accumulation of acetoin as well (up to 2.3 g/L) under optimized conditions (**Figure S4**). Glucose exhaustion in the supernatant at 48 h in both the optimized and control medium was confirmed by HPLC.

**Table 6.**
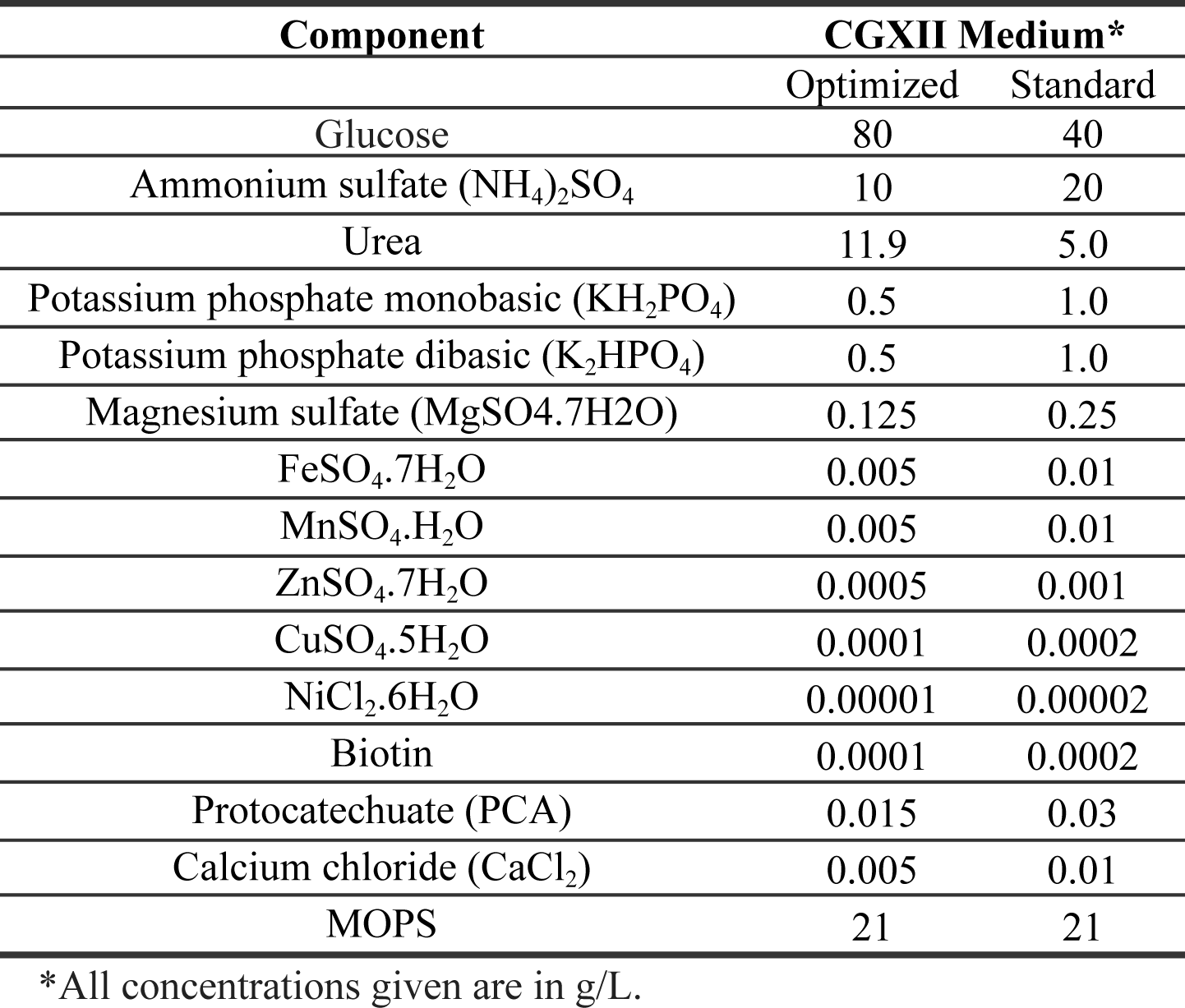
Concentrations of the Components Used in the Optimized and Standard CGXII Medium.

In the shake flask format, a drop in cell density compared to the control medium was observed. This might be due to more glucose being siphoned off for production of carotenoids (observed qualitatively as yellow cell pellets) in addition to TMP production compared to the control. In contrast to a published study (29) which shows optimal growth and production using a modified minimal medium with elimination of ammonium sulfate, we observed a significant drop in TMP titers (1650 mg/L) when no ammonium sulfate was used in the optimized media essentially reiterating the fact that ammonia is essential for the last spontaneous step in TMP production.

## 4. Conclusion

In this study, we have successfully engineered *C. glutamicum* to be a high titer production platform for TMP, establishing its robustness in real world carbon streams. Metabolic engineering and the optimization of the biosynthetic pathway allowed reliable production of TMP in both defined and complex media. We further optimized glucose based minimal medium components and conditions. We find that a high glucose concentration of 80 g/L and urea 11.9 g/L had the highest impact on productivity leading to up to 3.56 g/L TMP in 48 h of cultivation while the other minimal medium was reduced by half its original concentration in the CGXII medium. Functional genomics analysis would further help to shed light about competing pathways and candidates for targeted engineering and further conversion of accumulated acetoin to TMP using a two-stage process would also improve production. *C. glutamicum* is an important industrial host, and this study establishes it as a robust and stable production host for pyrazines and its precursors, which are commodity chemicals for a wide range of applications.

## Declarations

### Ethics Approval and Consent to Participate

Not Applicable

### Consent for Publication

All authors have read, provided feedback, and approved the manuscript for publication.

### Authors’ Contributions

Concept and Design: AM, AS, TE. Strain engineering and Media optimization: AS, KC-X. Metabolic model simulations: AS, DB. Generation of engineered Sorghum lines and hydrolysates: VP, AE, AO. Data analysis and Interpretation: AS, TE, AM. Supervision: AM, TE. Acquisition of funds: BAS, AM. Drafted the manuscript: AS, TE, AM.

## Acknowledgments

The authors would like to thank Dr. Mood Mohan, Postdoc affiliate, Lawrence Berkeley National Lab for his valuable suggestions for the statistical design of experiments.

## Data Availability

Data (including the additional files) is available upon request.

